# Industrializing yeast as a drug repurposing platform for inherited metabolic diseases

**DOI:** 10.1101/2024.08.23.609415

**Authors:** Mathuravani A. Thevandavakkam, Natalie E. Long, Briannna M. Roel, Kristin A. Kantautas, Shiri Zakin, Van Duesterberg, Ethan O. Perlstein

## Abstract

The development of therapies for rare diseases, particularly inherited metabolic disorders (IMDs), faces significant challenges due to the high cost and lengthy timelines involved. This study presents a yeast-based platform for drug repurposing that capitalizes on the remarkable similarity between yeast and human cellular pathways. This platform enables rapid, cost-effective screening of potential therapeutic compounds for rare diseases, offering a quick turnaround compared to traditional drug development processes. Utilizing a TargetMol library of comprising ∼50% nutraceuticals, our pipeline accelerates translation of promising drug repurposing hits into patient observational studies in as little as 6 months. We demonstrate the efficacy of this platform through three case studies in the context of IMDs, showcasing its potential to uncover novel treatments and reduce the time and expense associated with bringing therapies to patients with rare diseases.

## Introduction

Rare diseases, often of genetic origin, are defined as diseases that affect a small percentage of the population. With a demographically dependent definition of either affecting 1 in 200,000 individuals or with an occurrence rate of 1 in 2000, 95% of rare diseases lack an identified therapeutic approach [1,2] Collectively, there are nearly ten thousand rare diseases (and counting) with no treatment available. Among these are inherited metabolic diseases (IMDs), which are caused by deficiencies in metabolic enzymes resulting in imbalances in metabolic networks. With the rapid developments in whole genome sequencing offering the capabilities to diagnose new diseases sometimes as early as in infancy, the total number of rare (in particular ultra-rare) diseases is only expected to increase [3–5].

Rare diseases can manifest at any stage in life and are often chronic, progressive and incapacitating, resulting in substantial morbidity and mortality [6,7]. IMDs in particular manifest in neonates or early childhood and often present with significant cognitive impairments, neuromuscular degeneration, and multi-organ disturbances [8,9]. The diverse and individually uncommon nature of rare diseases, combined with the small population size they affect, presents a significant challenge in research and drug development and is financially unviable for the vast majority of pharmaceutical companies [10–13].

Perlara’s seminal work, “Yeast Models of Phosphomannomutase 2 Deficiency, a Congenital Disorder of Glycosylation” [14], laid the foundation for using a yeast CDG (Congenital Disorder of Glycosylation) disease model to screen small molecules to identify potential therapeutic targets and clinically actionable options for PMM2-CDG. The absence of other viable and phenotypically verifiable higher model systems led to the drug repurposing study in yeast that translated across PMM2-CDG models developed later. This led to the development of epalrestat – a generic diabetic neuropathy drug originally developed in Japan in 1990s – which is now being evaluated in a pivotal Phase III trial [15,16].

Drug repurposing, the process of identifying new therapeutic applications for existing drugs that have a well-established safety profile, offers a low cost, quick turnaround alternative for developing treatments for rare diseases, hastening drug development timelines with substantially reduced costs and risks (reviewed in [17]).

On-target and off-target drug repurposing efforts have identified modifiers of various disease phenotypes in late-onset neurodegenerative disorders such as Alzheimer’s, Parkinson’s diseases [18,19]. Currently, accessible treatments for these diseases do not fully ameliorate symptoms nor halt the progression of disease. However, the drugs most amenable for symptomatic management of these diseases were identified via drug repurposing approaches and include oncogenic kinase inhibitors and calcineurin inhibitors for Parkinson’s and Alzheimer’s disease, respectively. A compilation of drugs identified through drug repurposing illustrating its application for common late-onset neurodegenerative diseases is reviewed here [20,21]. Similar efforts in rare neurodegenerative disorders such as Wolfram syndrome using the iPSC model of the disease yielded repurposed drugs sodium valproate and dantrium [22].

Further, a high-throughput screen with >5,500 compounds in reprogrammed neurons identified sildenafil as a candidate therapeutic for mitochondrial DNA-associated Leigh syndrome (MILS) [23]. Sildenafil, an FDA-approved drug for pediatric pulmonary hypertension, demonstrated efficacy in enhancing intracellular calcium homeostasis and promoting neuronal growth in patient-derived cells [24,25]. Further, individual therapies in five patients showed that sildenafil was well tolerated and benefited Leigh syndrome patients resulting in sildenafil receiving a patent (EP 4 154 888 A1) and Orphan Drug Designation in Europe (EMA/OD/0000135459) highlighting its potential for repurposing in mitochondrial rare diseases.

Because of the evolutionary conservation between yeast and higher eukaryotes across many essential cellular processes not limited to DNA replication, transcription, and energy metabolism, yeast has served as a powerful model organism for studying human diseases and identifying potential therapeutic targets, enabling insights that are directly translatable to human biology [26–28].

Here we show that the well-established high-throughput screening platform using the eukaryotic model, *S*. *cerevisiae*, can effectively identify clinically actionable repurposed drugs for IMDs. This yeast-based platform is low-cost, rapid, scalable and most importantly leads to verifiable clinical outcomes via family-initiated and physician-guided “small N” observational studies.

## Results

### *S*. *cerevisiae* as disease avatar and a robust platform for high-throughput drug repurposing screens

To design a platform to enable modeling and screening for rare diseases, we first selected yeast strains that were orthologous and representative of the rare disease under study. The yeast genes were initially scored against human genes on percentage similarity, conservation of mutant residues, and functional ontology. Preliminary results suggested that yeast deletion strains (non-essential genes) or temperature-sensitive strains (essential genes) were equally informative in downstream screening and validation *versus* personalized yeast mutants representative of patient-specific variants (*unpublished data*). The use of commonly available mutant yeast strains, or yeast avatars, accelerated the clinical translation of compounds identified in a screen where cell growth is the unbiased phenotypic readout.

The screening platform described in Figure 1 comprises 3 phases: (1) phenotypic characterization of rare disease yeast avatars and their optimization; (2) a drug repurposing screen using the 2177 compound Pharmakon collection and/or the 8,387 compound L4000 Bioactives TargetMol library; (3) dose-response assays to validate the top rescuers (hits) and sensitizers from the TargetMol screen.

**Fig 1:**
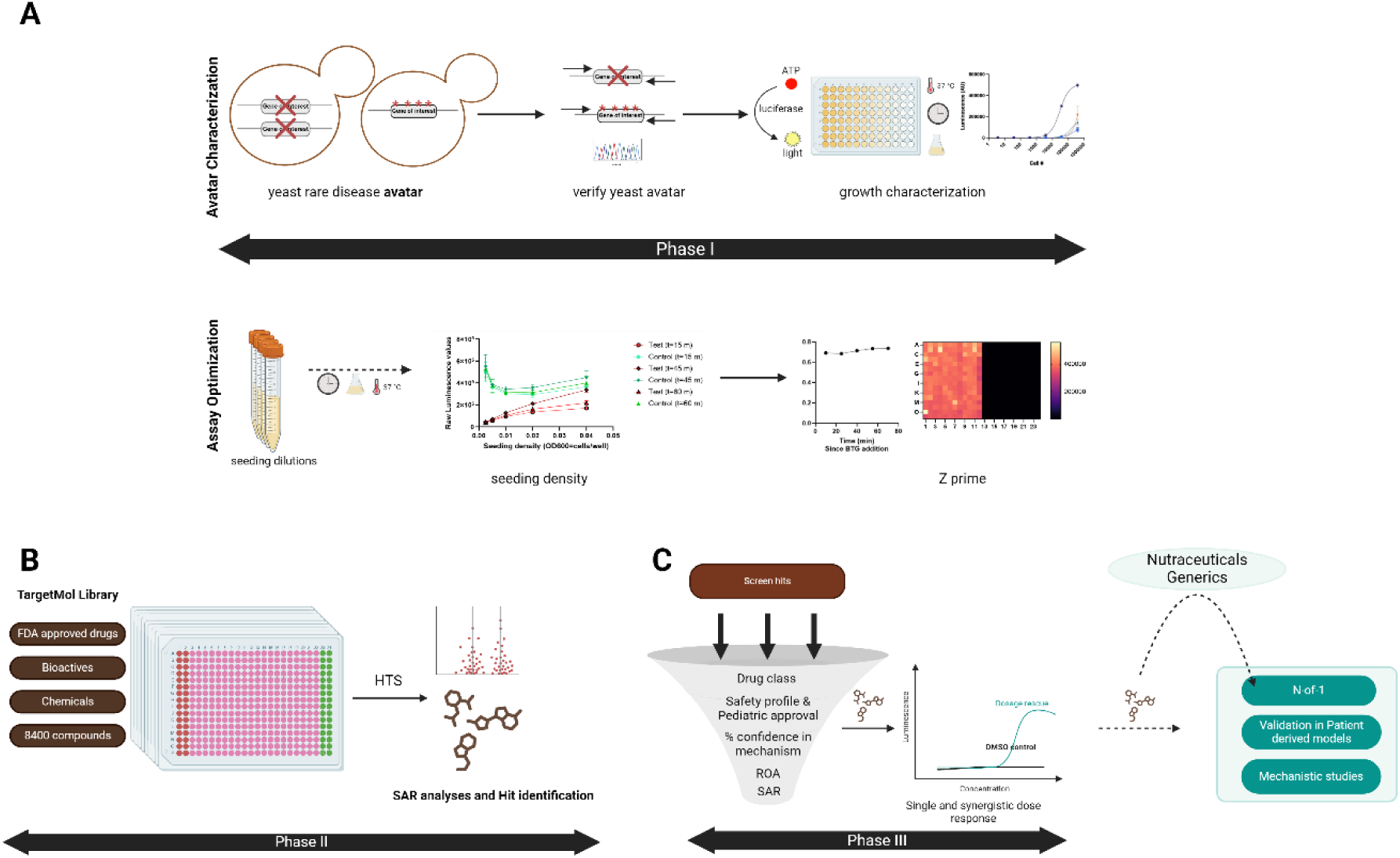
Yeast drug-repurposing workflow. The figure illustrates the workflow of the yeast drug repurposing screen. (A) Yeast homolog representative of a rare metabolic disease is procured from the yeast deletion/temperature-sensitive allele collection. The avatar is evaluated for growth deficiency and sensitized where required. Optimization experiments determine the ideal seeding density and Bac-titre Glo kinetics, with a Z’ score exceeding 0.5 being established before progressing the avatar to the screen. (B) The luminescence-based chemical screen is carried out using a pre-spotted TargetMol or Pharmakon library at a standard concentration of 20µM. Hits from the screen are scored based on their Z’ using the mean ± 3σ as cutoff. (C)The hits are classified on the mechanism of action and structure-and-activity relationship. Select hits undergo further hit validation in yeast (dose response assay/synergistic hit assessment) or patient derived cell lines (typically fibroblasts). Nutraceuticals are recommended for direct single patient observational study. (Image created with <u>BioRender</u>).

Yeast avatars for 16 rare diseases (Table 1) were characterized by growth at ambient and heat-sensitive conditions and under aerobic and anaerobic growth conditions where applicable. In contrast to traditional absorbance-based assays, we implemented a highly sensitive luminescence-based assay that measured the growth-deficient phenotype of yeast avatars compared to control strains (Fig 1a). Once the growth-deficient phenotype was established, optimization experiments were conducted to establish the optimal seeding density of cells, incubation time for optimal luminescence, and duration of growth for the screen (Fig 1b). A library of FDA and internationally approved drugs and nutraceuticals, or an FDA-approved library of repurposable drugs, was used in a one-shot (single-replicate) high-throughput screen. Compounds from the screen were scored as hits and sensitizers (see Methods). The top rescuers and sensitizers from the screen were validated in an 8-point dose-response assay, in triplicate. Select compounds were advanced for further validation in patient-derived disease models.

**Table 1:**
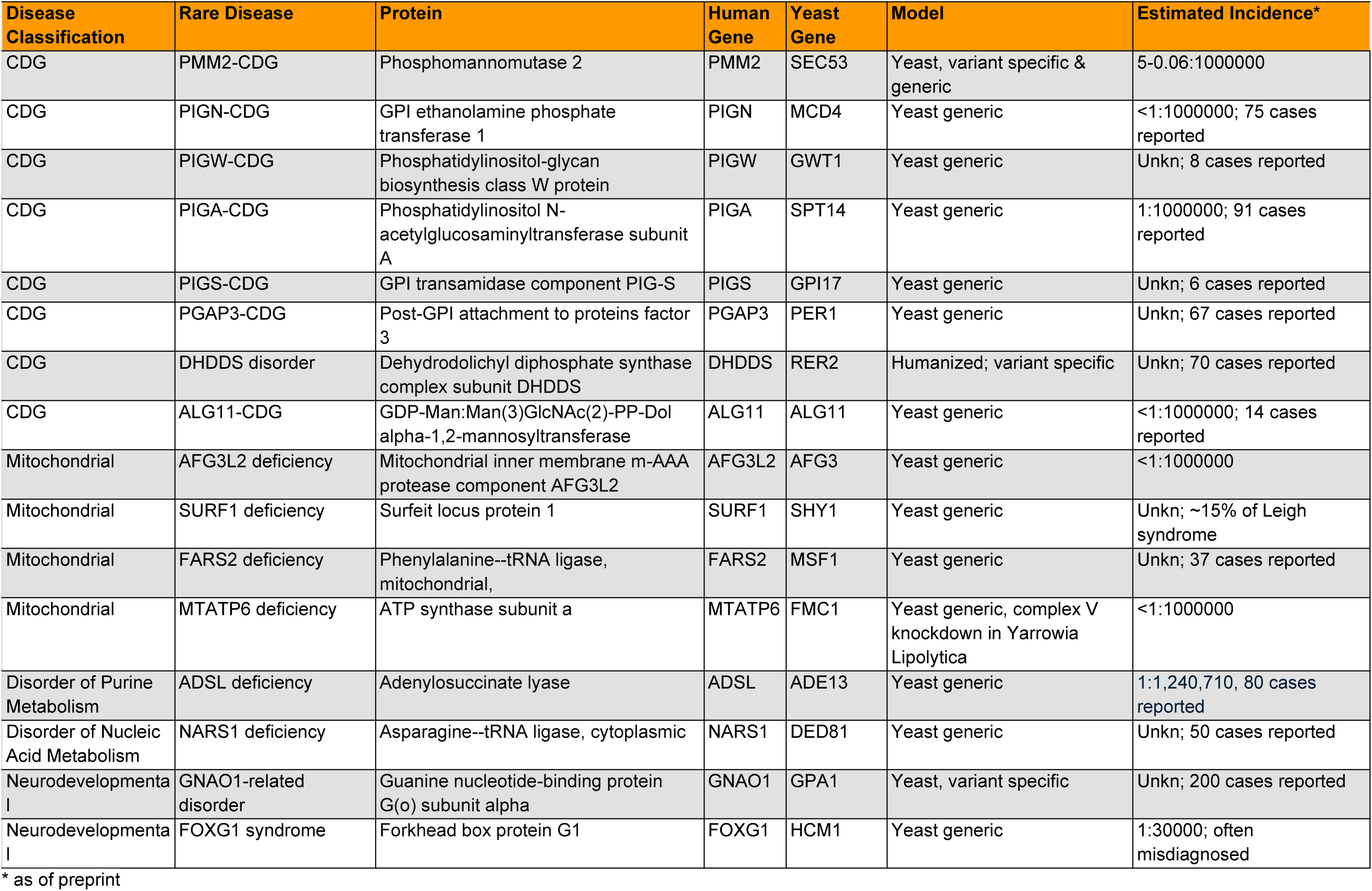
IMDs modeled in yeast in this study.

### A *S. cerevisiae* Drug Repurposing Platform for Targeting Mitochondrial, Congenital Disorders of Glycosylation and other Rare Inherited Metabolic Diseases

As shown in Table 1, the majority of Perlara’s yeast drug repurposing pipeline comprises inherited metabolic diseases (IMDs), particularly mitochondrial and CDG diseases. CDGs are a group of IMDs that disrupt the glycosylation process, which involves the addition of sugar molecules to proteins and lipids [29–31]. Defects in mitochondrial function, including disruptions in mitochondrial fusion, electron transport chain activity, ion transport, or organelle quality control, potentially lead to rare and ultra-rare conditions [32,33]. In both groups of diseases, patients often exhibit multi-systemic symptoms, including developmental delay, failure to thrive, hypotonia, neurological abnormalities, digestive and liver issues that may be accompanied by eye, skin, and heart conditions.

The IMDs under investigation are linked to loss-of-function mutations that reduce the activity of specific proteins. The mutant yeast strains used herein mimic the diminished or absent protein function seen in patients. The yeast avatars are subjected to conditions optimized for screening, allowing us to study the consequences of protein dysfunction in a controlled environment. For each yeast avatar examined (Fig 2), at least one lead compound was identified. This resulted in either a direct 1-to-N observational study (starting with a single patient and then incrementally scaling up with additional patients) or additional validation in patient-derived cell models, offering further mechanistic insights into disease biology. A detailed discussion of the hits or disease biology for each disease model is outside the scope of this paper.

**Fig 2:**
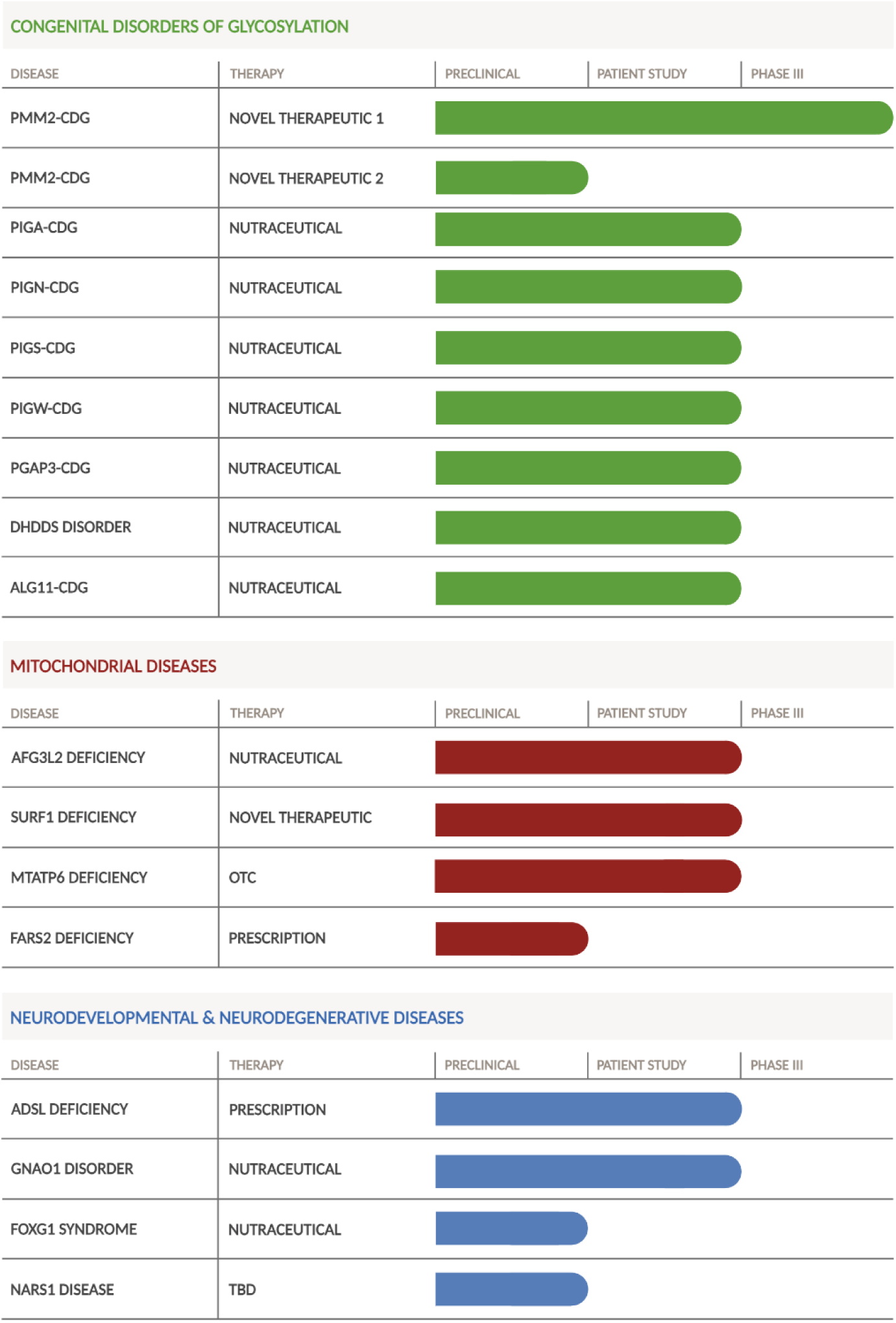
Perlara’s Yeast Drug-Repurposing pipeline. An overview of Perlara’s yeast drug repurposing pipeline and the progress made thus far, from early discovery and preclinical development to clinical trials across rare diseases. Key stages include drug identification, preclinical validation, and patient study and phased clinical trial, highlighting progress and milestones for each stage. (Image created with <u>BioRender</u>).

### Case study 1: mitochondrial deficiencies AFG3L2 and FARS2

We focused on two genes responsible for rare mitochondrial diseases: (1) AFG3L2— associated with spinocerebellar ataxia type 28 (SCA28), a neurodegenerative disorder characterized by progressive ataxia, dysarthria, and gait disturbances. This condition primarily affects mitochondrial protein quality control, with AFG3L2 mutations disrupting mitochondrial dynamics and function [34,35], and (2) FARS2—implicated in mitochondrial disease with a broader spectrum of clinical manifestations, including neurodevelopmental disorders and early-onset neurodegeneration. FARS2 mutations affect mitochondrial translation, leading to defective mitochondrial protein synthesis and impaired cellular energy production [36–38].

Both of these genes are conserved as 1:1 orthologs in S. *c*e*revisiae*. Using the yeast drug repurposing platform described above, under aerobic conditions to maximize mitochondrial dependency and thereby physiological dysfunction, we conducted an ∼8,400-compound TargetMol screen to identify potential therapeutic compounds that rescue mutant cell growth (Fig 3). Both screens displayed a tight clustering of data points around the mean, indicating robust assay performance and confidence in the results.

**Fig 3:**
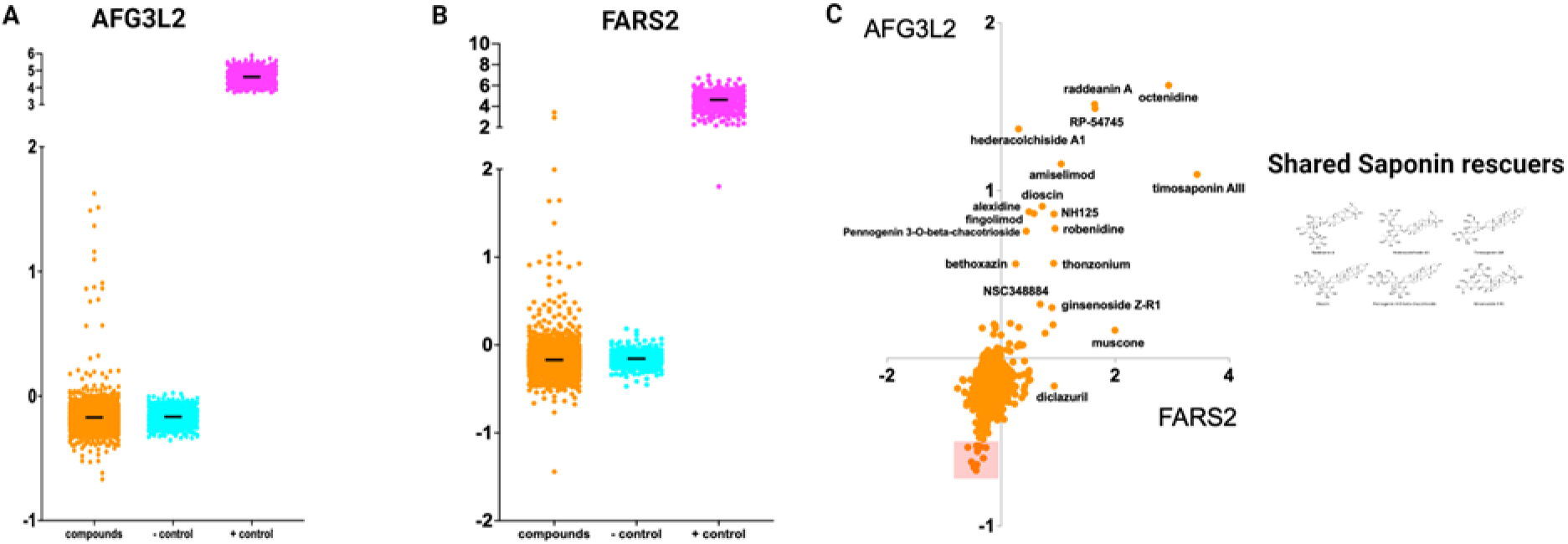
Comparison of TargetMol screen results for AFG3L2 and FARS2 mitochondrial deficiencies. (A) & (B) Summary plot of 8400 compound TargetMol screen in YP 2% lactate. Orange dots represent individual test compounds and the AFG3L2 and FARS2 knockout yeast avatar respectively. Cyan dots represent the negative control, AFG3L2 and FARS2 knockout yeast avatar treated with DMSO respectively, while the positive control, wildtype yeast treated with DMSO, is depicted as magenta dots. Despite some variability in wildtype yeast growth in the FARS2 screen, exceptionally tight distribution was observed in the negative controls. (C) Scatter plots categorize compounds as enhancers or sensitizers across both screens. The upper right quadrant is populated with shared rescuers while the lower left quadrant is empty, indicating the absence of shared sensitizers.

Intriguingly, the same class of compounds — steroidal saponins, quaternary amine-containing lipids, and sphingolipid analogs — effective for AFG3L2, also showed efficacy in FARS2 despite different underlying disease mechanisms. This convergence suggests that these compounds may alter mitochondrial membrane properties via alteration of lipid ratios, e.g., the ratio of phosphatidylcholine to phosphatidylethanolamine, enhancing the cell’s ability to manage proteotoxic stress. Further comparison of the AFG3L2 and FARS2 TargetMol Z score datasets revealed additional shared rescuers, including the quaternary amine lipid octenidine and the immunomodulating compound RP-54745, reinforcing the hypothesis of a common rescue mechanism for these mitochondrial diseases. It is not expected that growth in these screens would be rescued to wild-type levels consistent with complex interactions and mechanisms in play (see Discussion). Further, sensitizers of AFG3L2 and FARS2 (which were filtered to remove those toxic to all yeast avatars across screens) did not overlap. Although these diseases were all mitochondrial in nature, each exhibited distinct, disease-specific cellular responses.

### Case study 2: PIGA-CDG and PIGS-CDG

The PIGS protein, a component of the GPI transamidase (GPI-TA) complex, is crucial for attaching proteins to the GPI anchor. Dysfunction in PIGS can lead to the accumulation of unattached GPI anchors in the ER membrane [39]. To investigate potential rescue mechanisms, we utilized a *gpi17* temperature-sensitive (ts) yeast mutant that mimics PIGS deficiency. Upon exposure to 38°C, the mutant yeast exhibited a growth defect as expected.

We screened the Pharmakon library and identified over a dozen rescuers and several dozen sensitizers for PIGS (Fig 4a). Across all the Pharmakon screens analyzed, the SURF1 haploid and SURF1 diploid screens exhibited consistent patterns of rescuers and sensitizers, as expected, which bolstered confidence in the identified hits. The PGAP3 screen, however, displayed considerable noise — likely due to it being one of the initial screens conducted and the fact that PGAP3 deficiency in yeast is not lethal. Assay optimization was subsequently refined, including limiting the BTG incubation time to less than one hour, in order to reduce the signal-to-noise ratio. The PIGA, PIGS, ALG11, and ADSL screens predominantly identified sensitizers, with PIGA and ADSL screens notably yielding few rescuers.

**Fig 4:**
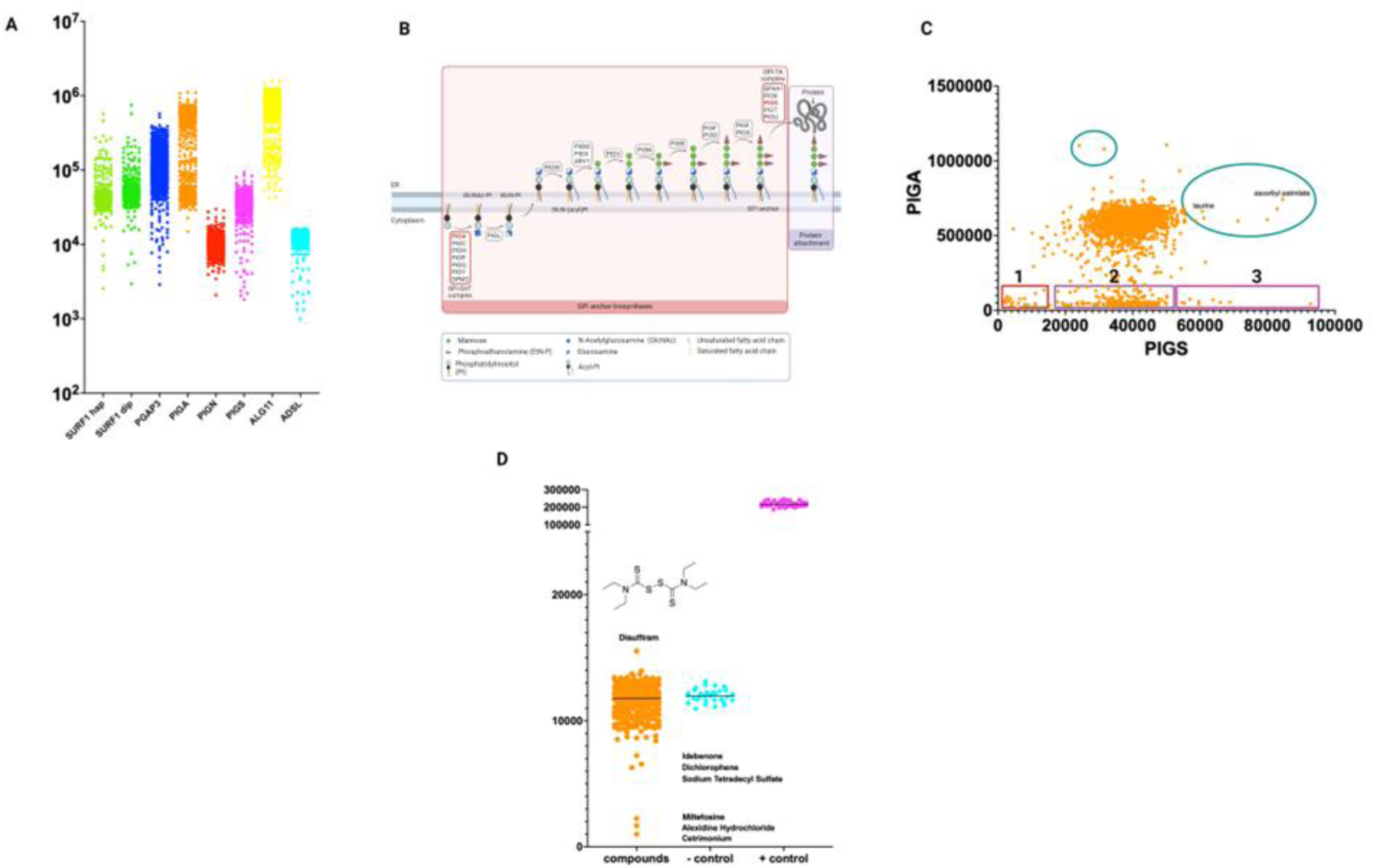
Comparative Analysis of Pharmakon Screens. (A) Summary plot of the 2177 compound Pharmakon screen. (B) Schematic of the glycosylphosphatidylinositol (GPI) anchor biosynthesis pathway, indicating key enzymes and their respective steps. (C) Scatter plot illustrating the relationship between PIGA and PIGS rescuers and sensitizers across various compounds. Sensitizers identified in the PIGA screen are categorized into three groups: (1) common sensitizers affecting both PIGA and PIGS, (2) PIGA sensitizers that partially rescue PIGS, and (3) PIGA sensitizers that significantly rescue PIGS. Green circles represent rescuers for PIGS and PIGA that do not overlap, indicating unique effects on each target. (D) Summary plot of single plate of Pharmakon showing disulfiram as the top rescuer for ADSL.

We compared the PIGS screen with the PIGA Pharmakon screen, as PIGA and PIGS function at opposite ends of the GPI anchor pathway. PIGA, as part of the GPI GlcNAc transferase (GPI-GnT) complex, catalyzes the transfer of GlcNAc to the lipid carrier molecule, forming an essential intermediate for the subsequent steps in GPI anchor synthesis [40]. PIGS is involved in GPI anchor attachment to proteins (Fig 4b).Compounds could be segregated into three sensitizer groups: those that were common sensitizers of both PIGA and PIGS; compounds that were unique sensitizers of either PIGA or PIGS; and, interestingly, compounds that sensitized PIGA but rescued PIGS (Fig 4c). There was little overlap between PIGA and PIGS rescuers. While the PIGS screen revealed a balance between sensitizers and rescuers, the PIGA screen was skewed toward identifying more sensitizers. This difference likely arises because the PIGA gene operates at the very first step of GPI anchor biosynthesis, a critical point that initiates the entire pathway and sets the stage for subsequent reactions. Disruption at this early stage can create a bottleneck, where the reduced production of functional GPI anchors affects the attachment and function of multiple downstream proteins. Modulation of PIGA activity can thus significantly alter GPI anchor production, leading to either a shortage or excess of intermediates, which disrupts cellular homeostasis and triggers more pronounced stress or dysfunction. As a result, disruptions at this stage are more deleterious compared to modulations of downstream proteins like PIGS.

Among the rescuers, ascorbyl palmitate (a fat-soluble form of Vitamin C) stood out as a potent rescuer for PIGS, although its mechanism of action remains unclear. Ascorbyl palmitate is currently being evaluated in a single PIGS-CDG patient observational study and further studies are planned to validate these findings in a PIGS-CDG fly model.

### Case study 3: Disulfiram’s Potential for ADSL Deficiency

The ADSL gene encodes adenylosuccinate lyase, an enzyme involved in two key steps of the purine biosynthesis pathway. ADSL catalyzes the conversion of succinylaminoimidazole carboxamide ribotide (SAICAR) to aminoimidazole carboxamide ribotide (AICAR) and the conversion of adenylosuccinate (S-AMP) to adenosine monophosphate (AMP). Deficiency in ADSL leads to the accumulation of toxic metabolites such as SAICAR and succinyladenosine (S-Ado), while reducing the levels of essential metabolites like fumarate [41,42]. ADSL deficiency was modeled using *ade13* temperature-sensitive mutant yeast strain.

From the Pharmakon screen, we identified seven effective rescuers and 27 sensitizers for ADSL deficiency. Notably, disulfiram emerged as a prominent rescuer. This finding is significant as disulfiram, typically a sensitizer in other screens, demonstrated a rescuing effect in the ADSL yeast model (Fig 4d). This result is unprecedented and suggests that disulfiram’s effects are context-dependent. Disulfiram, originally used for alcohol cessation, targets aldehyde dehydrogenase, but its mechanism of action in rescuing ADSL-deficient yeast remains unclear. Notably, a 2019 Chinese patent describes a novel use of disulfiram for treating ADSL deficiency, highlighting a potential link that warrants further investigation (CN111698986A). Preliminary studies in a worm model of ADSL recapitulates the rescue observed in the yeast ADSL model with disulfiram being evaluated in a single patient observational study.

Overall, our successful yeast modeling and identification of novel drug candidates highlight the potential for repurposing existing drugs and underscore the utility of yeast models in advancing therapeutic research for rare diseases.

### Yeast screens generate novel mechanistic insights into disease biology

A multi-screen meta-analysis revealed a significant overlap between hits identified in mitochondrial screens and those in CDG screens. This overlap suggests shared pathways or mechanisms affected by these hits, indicating potential common therapeutic targets for both types of diseases. The SURF1 and FARS2 yeast avatars have an overlapping distribution of hits, as expected in a comparison between two mitochondrial deficiencies. By contrast, a comparison of the TargetMol screening results using PMM2-CDG as a representative CDG and SURF1 as a representative mitochondrial deficiency reveals that the rescuing hits identified in mitochondrial screens exacerbate the CDG phenotype, but the converse, i.e., CDG rescuing hits worsening the mitochondrial phenotype, wasn’t always true (Fig 5a). These reciprocal effects highlight the intricate relationship between mitochondrial dysfunction and CDG, emphasizing the need for a holistic approach when developing therapeutic strategies for IMDs. Interestingly, mitochondrial dysfunction has been reported in several CDG types [16,43–46].

**Fig 5:**
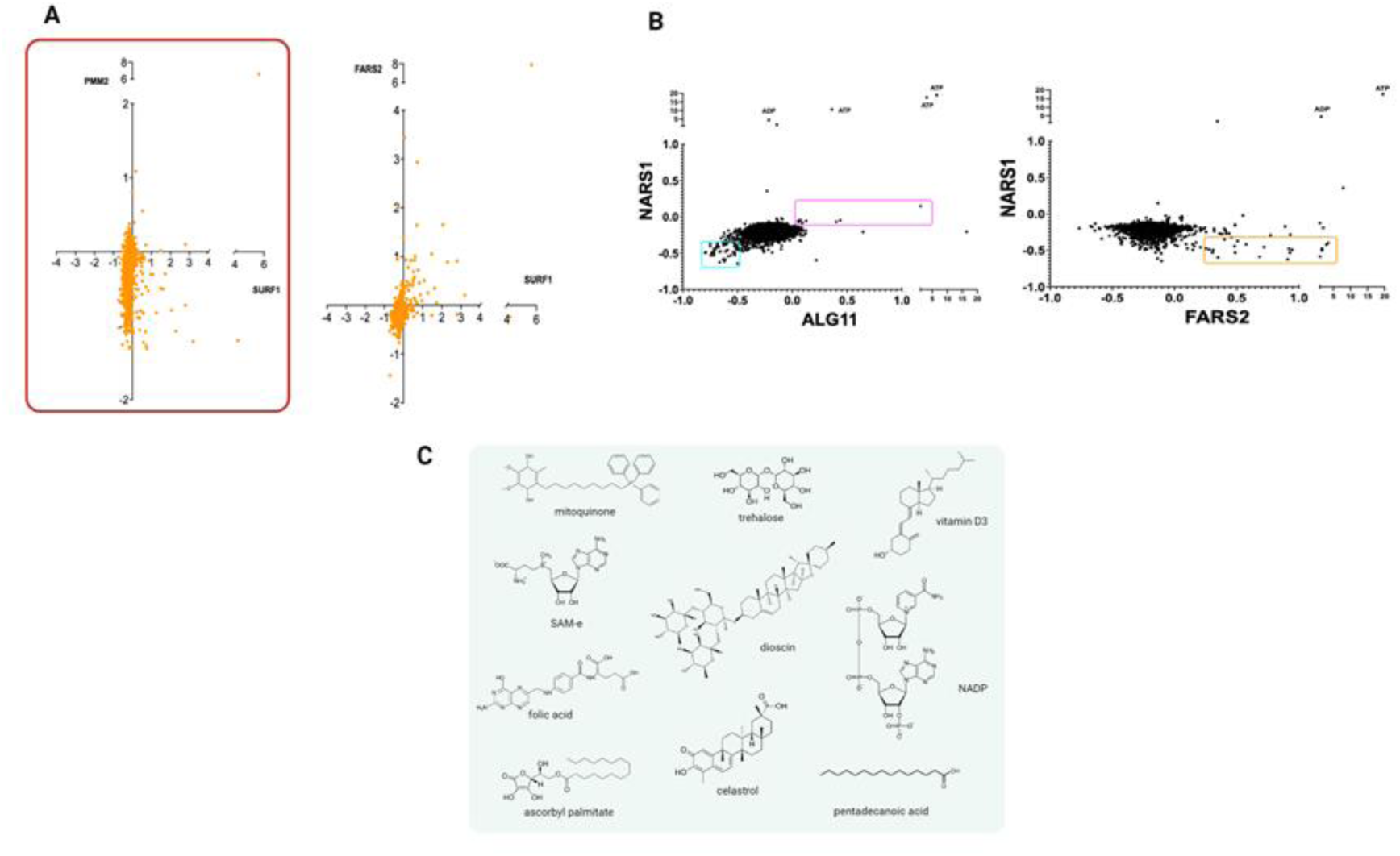
Patterns and mechanistic insights gained from yeast rare disease screens. (A) Scatter plot comparing hits and sensitizers across a CDG screen (PMM2) and mitochondrial screen (SURF1). Typically, compounds that rescue mitochondrial disease phenotype sensitize CDG yeast avatars (lower right quadrant), but not vice versa. (B) Scatter plots comparing NARS1 (cytoplasmic asparagine tRNA synthetase) and ALG11-CDG reveal overlap in sensitizers, while those NARS1 and FARS2 (a mitochondrial phenylalanine tRNA synthetase) are well-separated. (C) Examples of nutraceuticals: Examples of nutraceutical compounds that have been identified in TargetMol screens, demonstrating their relevance to drug repurposing efforts.

That observation led us to explore the relationship between NARS1 and CDGs. NARS1 (Asparaginyl-tRNA Synthetase 1) deficiency is associated with early-onset neurodevelopmental disorders and multi-organ involvement, differing from other aminoacyl-tRNA synthetase (ARS) deficiencies which typically present with progressive peripheral neuropathy later in life. Since NARS1 function is conserved in yeast, the yeast screening platform was leveraged to identify compounds that rescue the NARS1 deficiency. Strikingly, a comparison of this dataset to others revealed substantial overlap between NARS1 and ALG11 (a protein involved in early N-linked glycosylation that causes ALG11-CDG), with several shared rescuers and sensitizers, while the comparison between NARS1 and FARS2 (mitochondrial phenylalanine-tRNA Synthetase 2) showed minimal overlap. The sensitizers identified in NARS1 are consistent with those found in various CDG screens (Fig 5b).

Given that NARS1 is associated with early-onset neurodevelopmental delays and multi-organ disorders [47], it may be more accurately classified as a CDG rather than a Charcot-Marie-Tooth disorder. N-linked glycosylation, which involves attaching sugars to proteins via asparagine, could be disrupted in NARS1 deficiency. This would impact protein glycosylation due to decreased incorporation of asparagine residues into nascent proteins, leading to under-glycosylated or hypoglycosylated proteins. Asparagine levels may also be altered in patients, further linking NARS1 deficiency to CDG. The overlap of rescuers and sensitizers between NARS1 and CDGs, and the distinct clinical presentation, supports the classification of NARS1 deficiency as a type of CDG. This finding further underscores the importance of the vast screening dataset.

### Fast-tracking yeast nutraceutical hits to 1-of-N studies

In this study, using an expanded ∼8,400-compound TargetMol library of 50% nutraceutical compounds, we identified a clinically actionable nutraceutical as a rescuer in nearly every screen (Fig 5c). Often with an established pediatric safety profile, the regulatory ease, accessibility and cost-effectiveness of these compounds enable rapid translation of candidate medicines to the clinic. Our results underscore the untapped potential of nutraceuticals as therapeutic options for IMDs. Several nutraceuticals identified in the screens demonstrated promising effects in terms of their growth rescue effects, suggesting they could be explored further as novel treatments for rare and complex metabolic conditions. In about 50% of rare diseases where drug repurposing candidates were identified using the Perlara’s yeast pipeline, nutraceuticals emerged as a top hit. These hits are being validated in single patient observational studies, significantly accelerating the clinical translation timeline from decades to as little as 6 months.

We compared the dataset from our previous PMM2-CDG yeast-based screens (of several haploid and diploid PMM2 patient alleles [14], with the results from a drug repurposing screen using a compound heterozygous mutant yeast strain corresponding to a different PMM2 patient genotype. The same structure-activity relationships were identified in both screens (*unpublished data*). However, the use of a more comprehensive library, coupled to the increased sensitivity of our screening platform, revealed hits with novel mechanisms of action that are under further study.

Collectively, these studies support the efficacy of using off-the-shelf or designer mutant yeast strains as rare disease patient avatars, demonstrating assay consistency, data reliability, and clinical actionability across different experimental conditions. Overall, these findings contribute to a deeper understanding of the interactions between mitochondrial dysfunction and glycosylation disorders, reveal new therapeutic avenues, and accelerate new therapeutics development.

## Discussion

We developed a cost-effective, rapid-turnaround, high-throughput pipeline for drug repurposing in rare genetic metabolic diseases using *Saccharomyces cerevisiae*. This pipeline leverages readily available yeast deletion and temperature-sensitive allele collections for drug repurposing screens, facilitating the rapid clinical translation of therapeutic options for IMDs. Our luminescence-based approach significantly increased the sensitivity and reliability of the screens. By utilizing a comprehensive drug library that includes a balanced 50:50 ratio of nutraceuticals to approved drugs and generics, we enhanced the rapid and reliable translation of screen hits to 1-to-N observational studies in patients. We confirmed the reliability of our platform approach by comparing disease-specific avatar-based screens to those using a generic yeast mutant strain for the gene of interest. The efficacy of our pipeline is demonstrated by the number of compounds discovered through these screens that have been successfully validated in 1-to-N trials and in patient-derived model systems, including fibroblasts, neuronal progenitor cells, and organoids [48].

The deep evolutionary conservation between yeast and human genes, combined with using yeast cell growth as a single-variable readout of a multifactorial disease state, is the cornerstone of our drug-repurposing platform. Yeast, particularly *Saccharomyces cerevisiae*, shares a substantial number of homologous genes with humans, especially those involved in fundamental cellular processes such as metabolism, DNA replication, and protein synthesis. This evolutionary conservation extends to the network level, allowing us to model the cellular pathophysiology of human diseases in yeast with high fidelity, dissect the underlying mechanisms of these diseases, and identify disease modifiers [26–28]. We intentionally screen for compounds that restore balance to evolutionarily conserved pathways, increasing the likelihood that these compounds will have similar effects in human cells.

Perlara’s current portfolio is focused on IMDs, a large percentage of which include mitochondrial diseases and congenital disorders of glycosylation (CDGs). Two important constraints of studying such diseases have been assessed in our pipeline:

*Suitability of S.cerevisiae in modeling mitochondrial diseases: S. cerevisiae* lacks Complex I of the respiratory chain, unlike humans, which introduces complexity in modeling mitochondrial diseases. However, our screening pipeline addresses this gap by incorporating both *S. cerevisiae* and *Yarrowia lipolytica*. For instance, in MTATP6 Leigh syndrome, where mutations in MTATP6 impair ATP synthase (Complex V) function, we conducted parallel drug repurposing screens using *S. cerevisiae* (bearing FMC1 deletion under aerobic conditions) and *Y. lipolytica* (with Complex V disrupted by oligomycin). Analysis of hits and rescuers from these screens revealed both unique and conserved structure-activity relationships among compounds that mitigated the growth defect (*unpublished data*). The combined insights from both yeast models may alleviate disease pathology in MTATP6 Leigh syndrome.

Additionally, *Y. lipolytica* has been used successfully in drug repurposing screens for Complex I deficiency, as previously published by our group [49]. The ability to use any yeast species and integrate their screening results maximizes the information obtained and overcomes challenges associated with using yeast as a model system where whole pathways are not faithfully conserved. A dual-yeast-species approach enhances the robustness and applicability of our drug-repurposing platform for mitochondrial diseases. Further, repurposed drugs identified via the platform for SURF1 Leigh’s syndrome model were validated in midbrain organoids of Leigh syndrome where the compounds improved both neurogenesis and neurite organization underscoring the clinical translatability and reliability of yeast models of mitochondrial diseases [48].

*Suitability of S.cerevisiae in modeling defects in cholesterol biosynthesis:S. cerevisiae* serves as a powerful model due to the conservation of fundamental cellular processes. However, one significant difference between yeast and humans is the lack of cholesterol biosynthesis in yeast. Instead, yeast synthesizes ergosterol, a sterol with similar structural and functional roles in cellular membranes, sharing many enzymatic steps with the cholesterol biosynthetic pathway in humans [50,51]. This conservation allows for the study of sterol biosynthesis and its regulation using yeast as a model. Despite the difference in the end product (ergosterol vs. cholesterol), many enzymes involved in sterol biosynthesis are homologous between yeast and humans, providing a basis for understanding sterol metabolism and its regulation. This is particularly evident in CDGs, where there is an intricate link between glycosylation and lipid metabolism. Many CDGs can be modeled in yeast, and the expression of human genes in yeast can compensate for the lack of the yeast gene.

Yeast strains with humanized cholesterol biosynthesis [52] may be employed in the same pipeline in the future if current models do not fully encapsulate the range of defects observed in human diseases. However, our experience indicates that compounds altering ergosterol biosynthesis, such as sertaconazole in SURF1 Leigh disease models, exert similar effects on cholesterol biosynthesis in organoid models. Further ongoing validation in CDG disease models suggest that this difference does not compromise the translatability of hits from yeast to 1-to-N trials.

Our pipeline is designed to address the various challenges presented by human rare diseases and has been rigorously tested for its limitations, ensuring robust and translatable findings.

The short-term economic constraints of rare disease research and the long-term pharmacoeconomic costs of rare disease care necessitate cost-effective approaches that can be implemented as early in life as possible. Yeast-based drug repurposing is inherently low-cost due to the simplicity and affordability of yeast culture and genetic manipulation. This approach not only reduces the costs and barriers to entry of drug screening, but also promises a fast lane to clinical interventions that can be implemented immediately after diagnosis. Consequently, our platform has significant potential for discovering affordable and accessible life-changing treatments for rare disease communities starting with inherited metabolic diseases.

## Methods

### Strains and plasmids

All strains used herein are in the S288C background (excepting ALG11 knockout strain in W303 background). Strains were grown in Yeast extract, Peptone and Dextrose (YPD) media (RPI Research Product International) at 30°C unless otherwise noted. Standard procedures were followed for yeast maintenance (see Table 2 for list of strains and growth conditions)

**Table 2:**
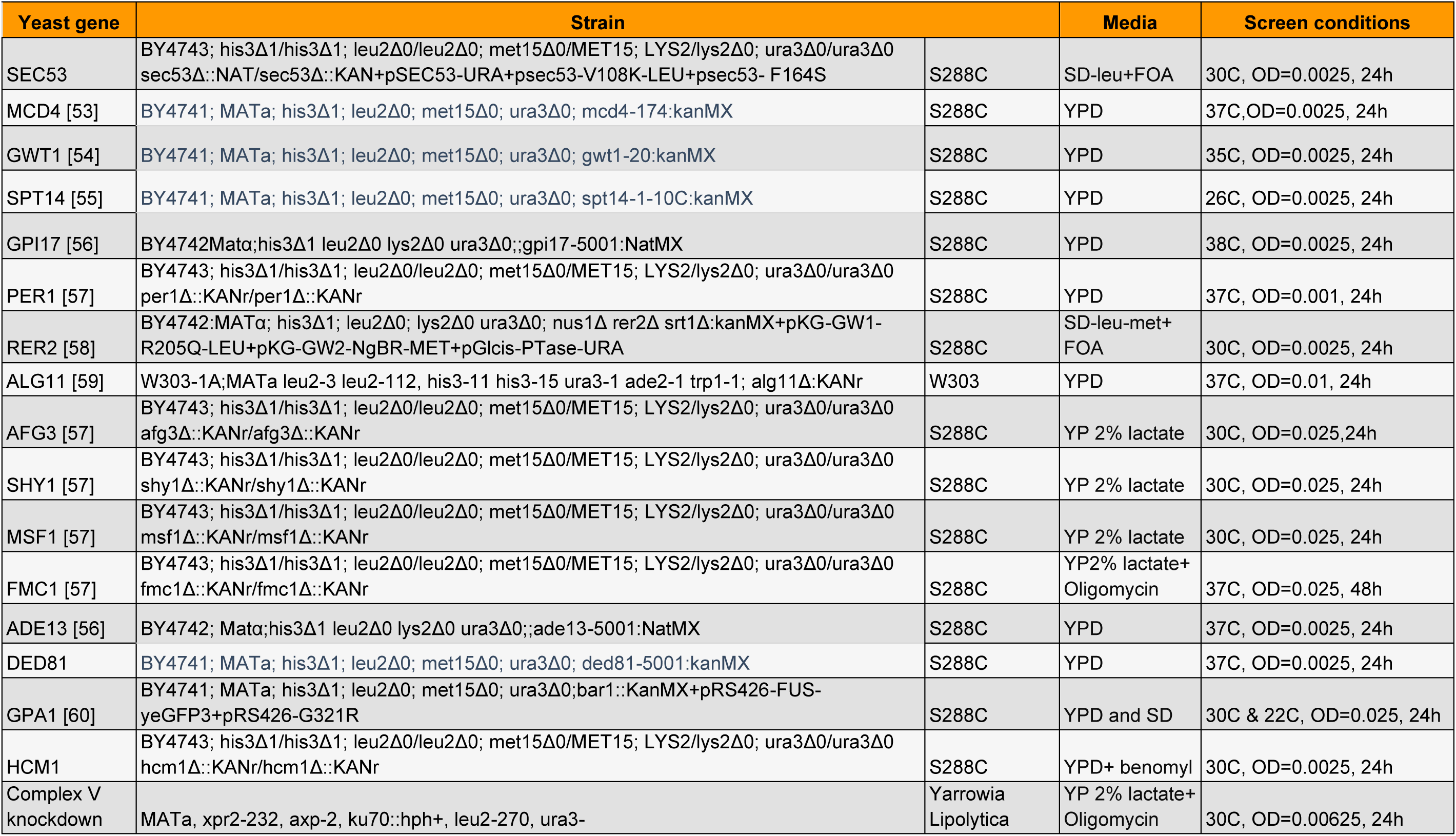
Yeast strains used in the study.

| Yeast gene | Strain |  | Media | Screen conditions |
| --- | --- | --- | --- | --- |
| SEC53 | BY4743; his3Δ1/his3Δ1; leu2Δ0/leu2Δ0; met15Δ0/MET15; LYS2/lys2Δ0; ura3Δ0/ura3Δ0<br>sec53Δ::NAT/sec53Δ::KAN+pSEC53-URA+psec53-V108K-LEU+psec53- F164S | S288C | SD-leu+FOA | 30C, OD=0.0025, 24h |
| MCD4 [53] | BY4741; MATa; his3Δ1; leu2Δ0; met15Δ0; ura3Δ0; mcd4-174:kanMX | S288C | YPD | 37C, OD=0.0025, 24h |
| GWT1 [54] | BY4741; MATa; his3Δ1; leu2Δ0; met15Δ0; ura3Δ0; gwt1-20:kanMX | S288C | YPD | 35C, OD=0.0025, 24h |
| SPT14 [55] | BY4741; MATa; his3Δ1; leu2Δ0; met15Δ0; ura3Δ0; spt14-1-10C:kanMX | S288C | YPD | 26C, OD=0.0025, 24h |
| GPI17 [56] | BY4742Matα;his3Δ1 leu2Δ0 lys2Δ0 ura3Δ0;;gpi17-5001:NatMX | S288C | YPD | 38C, OD=0.0025, 24h |
| PER1 [57] | BY4743; his3Δ1/his3Δ1; leu2Δ0/leu2Δ0; met15Δ0/MET15; LYS2/lys2Δ0; ura3Δ0/ura3Δ0<br>per1Δ::KANr/per1Δ::KANr | S288C | YPD | 37C, OD=0.001, 24h |
| RER2 [58] | BY4742:MATα; his3Δ1; leu2Δ0; lys2Δ0 ura3Δ0; nus1Δ rer2Δ srt1Δ:kanMX+pKG-GW1-<br>R205Q-LEU+pKG-GW2-NgBR-MET+pGlcis-PTase-URA | S288C | SD-leu-met+<br>FOA | 30C, OD=0.0025, 24h |
| ALG11 [59] | W303-1A;MATa leu2-3 leu2-112, his3-11 his3-15 ura3-1 ade2-1 trp1-1; alg11Δ:KANr | W303 | YPD | 37C, OD=0.01, 24h |
| AFG3 [57] | BY4743; his3Δ1/his3Δ1; leu2Δ0/leu2Δ0; met15Δ0/MET15; LYS2/lys2Δ0; ura3Δ0/ura3Δ0<br>afg3Δ::KANr/afg3Δ::KANr | S288C | YP 2% lactate | 30C, OD=0.025, 24h |
| SHY1 [57] | BY4743; his3Δ1/his3Δ1; leu2Δ0/leu2Δ0; met15Δ0/MET15; LYS2/lys2Δ0; ura3Δ0/ura3Δ0<br>shy1Δ::KANr/shy1Δ::KANr | S288C | YP 2% lactate | 30C, OD=0.025, 24h |
| MSF1 [57] | BY4743; his3Δ1/his3Δ1; leu2Δ0/leu2Δ0; met15Δ0/MET15; LYS2/lys2Δ0; ura3Δ0/ura3Δ0<br>msf1Δ::KANr/msf1Δ::KANr | S288C | YP 2% lactate | 30C, OD=0.025, 24h |
| FMC1 [57] | BY4743; his3Δ1/his3Δ1; leu2Δ0/leu2Δ0; met15Δ0/MET15; LYS2/lys2Δ0; ura3Δ0/ura3Δ0<br>fmc1Δ::KANr/fmc1Δ::KANr | S288C | YP2% lactate+<br>Oligomycin | 37C, OD=0.025, 48h |
| ADE13 [56] | BY4742; Matα;his3Δ1 leu2Δ0 lys2Δ0 ura3Δ0;;ade13-5001:NatMX | S288C | YPD | 37C, OD=0.0025, 24h |
| DED81 | BY4741; MATa; his3Δ1; leu2Δ0; met15Δ0; ura3Δ0; ded81-5001:kanMX | S288C | YPD | 37C, OD=0.0025, 24h |
| GPA1 [60] | BY4741; MATa; his3Δ1; leu2Δ0; met15Δ0; ura3Δ0;bar1::KanMX+pRS426-FUS-<br>yeGFP3+pRS426-G321R | S288C | YPD and SD | 30C & 22C, OD=0.025, 24h |
| HCM1 | BY4743; his3Δ1/his3Δ1; leu2Δ0/leu2Δ0; met15Δ0/MET15; LYS2/lys2Δ0; ura3Δ0/ura3Δ0<br>hcm1Δ::KANr/hcm1Δ::KANr | S288C | YPD+ benomyl | 30C, OD=0.0025, 24h |
| Complex V<br>knockdown | MATa, xpr2-232, axp-2, ku70::hph+, leu2-270, ura3- | Yarrowia<br>Lipolytica | YP 2% lactate+<br>Oligomycin | 30C, OD=0.00625, 24h |

### Growth assay

Cells from overnight cultures were resuspended in YPD media to OD_6_oo = 0.4, then serial diluted into 100 µL YPD media in 96-well plates at 10^-1^, 10^-2^, 10^-3^, 10^-4^ and 10 ^-5^. Plates were incubated at growth deficient temperature (see Table 2) for 24 hours. 50 µL of yeast cell suspension was transferred to a 96-well plate with 50 µL of BacTiter-Glo (Promega), briefly vortexed and incubated for 5 minutes. Luminescence readings were measured by a plate reader (ThermoFisher Varioskan Lux Multimode Microplate Reader).

### Drug screens

Drug screens were performed using a 2,177-compound collection (Small Molecule Discovery Center (“SMDC”) Drug collection), and/or the 8,387 L4000 Bioactive Compound library (TargetMol). 50 nL of compounds the from SMDC Drug collection, 200 nL of compounds from the L4100 Bioactive Compound library, or DMSO, were dispensed into 384-well plates using the Echo acoustic dispenser (Beckman Coulter Labcyte Echo 650) to achieve a final concentration of 20 µM. Cells from overnight cultures were resuspended in suitable media to optimized OD_6_oo and 25 µL (20 µL for TargetMol) of yeast cell suspensions were dispensed into the 384-well plates containing compounds or DMSO with a EL406 automated dispenser (Biotek). Plates were covered and incubated at optimized temperature for 24-48 hours. 25 µL (20 µL for TargetMol) of BacTiter-Glo was dispensed into the plates. Plates were briefly vortexed and then incubated for optimized time (10-50) minutes prior to reading (Envision Plate reader).

### Dose response assays

Dose response assays were performed with select compounds identified through the drug screens Each compound was dispensed at eight concentrations in triplicate, beginning with 40 µM and following a two-fold dilution (40 µM, 20 µM, 10 µM, 5 µM, etc.), while DMSO was administered at a single concentration of 0.72% to control wells. Compounds or DMSO were dispensed into 384-well plate using the Echo acoustic dispenser (Beckman Coulter Labcyte Echo 650). Within the plate, 16 wells were allocated for the negative control (MCD4 mutant DMSO) and 16 wells for the positive control (wild type yeast DMSO). Cells from overnight cultures were resuspended in appropriate media to an optimized OD_6_oo and 20 µL of yeast cell suspensions were dispensed into the 384-well plate containing compounds or DMSO with a EL406 automated dispenser (Biotek). Plates were covered and incubated at appropriate temperature for 24-48 hours. 20 µL of BacTiter-Glo was dispensed into the plates. Plates were briefly vortexed and then incubated for optimized time (10-50) minutes prior to reading (Envision Plate reader).

## References

1. Nguengang Wakap S, Lambert DM, Olry A, et al. Estimating cumulative point prevalence of rare diseases: analysis of the Orphanet database. Eur J Hum Genet. 2020;28(2):165–173. doi:10.1038/s41431-019-0508-0

2. Haendel M, Vasilevsky N, Unni D, et al. How many rare diseases are there? Nat Rev Drug Discov. 2020;19(2):77–78. doi:10.1038/d41573-019-00180-y

3. Wojcik MH, Bresnahan M, Del Rosario MC, Ojeda MM, Kritzer A, Fraiman YS. Rare diseases, common barriers: disparities in pediatric clinical genetics outcomes. Pediatr Res. 2023;93(1):110–117. doi:10.1038/s41390-022-02240-3

4. Meng M, Zhang YP. Impact of inborn errors of metabolism on admission in a neonatal intensive care unit: a 4-year report. J Pediatr Endocrinol Metab. 2013;26(7-8):689–693. doi:10.1515/jpem-2013-0021

5. Navarrete-Opazo AA, Singh M, Tisdale A, Cutillo CM, Garrison SR. Can you hear us now? The impact of health-care utilization by rare disease patients in the United States. Genet Med. 2021;23(11):2194–2201. doi:10.1038/s41436-021-01241-7

6. Ferreira CR. The burden of rare diseases. Am J Med Genet A. 2019;179(6):885–892. doi:10.1002/ajmg.a.61124

7. Mazzucato M, Visonà Dalla Pozza L, Minichiello C, Toto E, Vianello A, Facchin P. Estimating mortality in rare diseases using a population-based registry, 2002 through 2019. Orphanet J Rare Dis. 2023;18(1):362. doi:10.1186/s13023-023-02944-7

8. Romão A, Simon PEA, Góes JEC, et al. INITIAL CLINICAL PRESENTATION IN CASES OF INBORN ERRORS OF METABOLISM IN A REFERENCE CHILDREN’S HOSPITAL: STILL A DIAGNOSTIC CHALLENGE. Rev Paul Pediatr. 2017;35(3):258–264. doi:10.1590/1984-0462/;2017;35;3;00012

9. Ferreira CR, van Karnebeek CDM, Vockley J, Blau N. A proposed nosology of inborn errors of metabolism. Genet Med. 2019;21(1):102–106. doi:10.1038/s41436-018-0022-8

10. Li X, He J, He L, et al. Spectrum Analysis of Inherited Metabolic Disorders for Expanded Newborn Screening in a Central Chinese Population. Front Genet. 2021;12:763222. doi:10.3389/fgene.2021.763222

11. Teixeira C, Cordeiro C, Pinto C, Diogo L. Clinical Presentation of Inherited Metabolic Diseases in Newborns Hospitalised in an Intensive Care Unit. J Mother Child. 2023;27(1):55–63. doi:10.34763/jmotherandchild.20232701.d-23-00021

12. Couce ML, Baña A, Bóveda MD, Pérez-Muñuzuri A, Fernández-Lorenzo JR, Fraga JM. Inborn errors of metabolism in a neonatology unit: impact and long-term results. Pediatr Int. 2011;53(1):13–17. doi:10.1111/j.1442-200X.2010.03177.x

13. Bower A, Imbard A, Benoist JF, et al. Diagnostic contribution of metabolic workup for neonatal inherited metabolic disorders in the absence of expanded newborn screening. Sci Rep. 2019;9(1):14098. doi:10.1038/s41598-019-50518-0

14. Lao JP, DiPrimio N, Prangley M, Sam FS, Mast JD, Perlstein EO. Yeast Models of Phosphomannomutase 2 Deficiency, a Congenital Disorder of Glycosylation. G3 (Bethesda). 2019;9(2):413-423. doi:10.1534/g3.118.200934

15. Maggie’s Pearl L. Oral Epalrestat Therapy in Pediatric Subjects With PMM2-CDG. ClinicalTrials.gov Identifier: NCT04925960. Last Update Posted: February 13, 2023. Accessed January 5, 2024. https://classic.clinicaltrials.gov/ct2/show/NCT04925960

16. Iyer S, Sam FS, DiPrimio N, et al. Repurposing the aldose reductase inhibitor and diabetic neuropathy drug epalrestat for the congenital disorder of glycosylation PMM2-CDG. Dis Model Mech. 2019;12(11). doi:10.1242/dmm.040584

17. Kulkarni VS, Alagarsamy V, Solomon VR, Jose PA, Murugesan S. Drug Repurposing: An Effective Tool in Modern Drug Discovery. Russ J Bioorg Chem. 2023;49(2):157–166. doi:10.1134/S1068162023020139

18. Durães F, Pinto M, Sousa E. Old Drugs as New Treatments for Neurodegenerative Diseases. Pharmaceuticals (Basel). 2018;11(2). doi:10.3390/ph11020044

19. Kakoti BB, Bezbaruah R, Ahmed N. Therapeutic drug repositioning with special emphasis on neurodegenerative diseases: Threats and issues. Front Pharmacol. 2022;13:1007315. doi:10.3389/fphar.2022.1007315

20. Shah S, Dooms MM, Amaral-Garcia S, Igoillo-Esteve M. Current Drug Repurposing Strategies for Rare Neurodegenerative Disorders. Front Pharmacol. 2021;12:768023. doi:10.3389/fphar.2021.768023

21. Alrouji M, Al-Kuraishy HM, Al-Gareeb AI, et al. Cyclin-dependent kinase 5 (CDK5) inhibitors in Parkinson disease. J Cell Mol Med. 2024;28(11):e18412. doi:10.1111/jcmm.18412

22. Abreu D, Urano F. Current Landscape of Treatments for Wolfram Syndrome. Trends Pharmacol Sci. 2019;40(10):711–714. doi:10.1016/j.tips.2019.07.011

23. Lorenz C, Lesimple P, Bukowiecki R, et al. Human iPSC-Derived Neural Progenitors Are an Effective Drug Discovery Model for Neurological mtDNA Disorders. Cell Stem Cell. 2017;20(5):659–674.e9. doi:10.1016/j.stem.2016.12.013

24. Barst RJ, Ivy DD, Gaitan G, et al. A randomized, double-blind, placebo-controlled, dose-ranging study of oral sildenafil citrate in treatment-naive children with pulmonary arterial hypertension. Circulation. 2012;125(2):324–334. doi:10.1161/CIRCULATIONAHA.110.016667

25. Humpl T, Reyes JT, Holtby H, Stephens D, Adatia I. Beneficial effect of oral sildenafil therapy on childhood pulmonary arterial hypertension: twelve-month clinical trial of a single-drug, open-label, pilot study. Circulation. 2005;111(24):3274–3280. doi:10.1161/CIRCULATIONAHA.104.473371

26. Botstein D, Fink GR. Yeast: an experimental organism for modern biology. Science. 1988;240(4858):1439–1443. doi:10.1126/science.3287619

27. Kachroo AH, Laurent JM, Yellman CM, Meyer AG, Wilke CO, Marcotte EM. Evolution. Systematic humanization of yeast genes reveals conserved functions and genetic modularity. Science. 2015;348(6237):921–925. doi:10.1126/science.aaa0769

28. Liu W, Li L, Ye H, et al. From Saccharomyces cerevisiae to human: The important gene co-expression modules. Biomed Rep. 2017;7(2):153–158. doi:10.3892/br.2017.941

29. Ondruskova N, Cechova A, Hansikova H, Honzik T, Jaeken J. Congenital disorders of glycosylation: Still “hot” in 2020. Biochim Biophys Acta Gen Subj. 2021;1865(1):129751. doi:10.1016/j.bbagen.2020.129751

30. Raynor A, Haouari W, Lebredonchel E, Foulquier F, Fenaille F, Bruneel A. Biochemical diagnosis of congenital disorders of glycosylation. Adv Clin Chem. 2024;120:1–43. doi:10.1016/bs.acc.2024.03.001

31. Gardeitchik T, Wyckmans J, Morava E. Complex Phenotypes in Inborn Errors of Metabolism: Overlapping Presentations in Congenital Disorders of Glycosylation and Mitochondrial Disorders. Pediatr Clin North Am. 2018;65(2):375–388. doi:10.1016/j.pcl.2017.11.012

32. Gorman GS, Chinnery PF, DiMauro S, et al. Mitochondrial diseases. Nat Rev Dis Primers. 2016;2. doi:10.1038/nrdp.2016.80

33. Barca E, Long Y, Cooley V, et al. Mitochondrial diseases in North America: An analysis of the NAMDC Registry. Neurol Genet. 2020;6(2). doi:10.1212/NXG.0000000000000402

34. Patron M, Sprenger HG, Langer T. m-AAA proteases, mitochondrial calcium homeostasis and neurodegeneration. Cell Res. 2018;28(3):296–306. doi:10.1038/cr.2018.17

35. Almajan ER, Richter R, Paeger L, et al. AFG3L2 supports mitochondrial protein synthesis and Purkinje cell survival. J Clin Invest. 2012;122(11):4048–4058. doi:10.1172/JCI64604

36. Chen W, Rehsi P, Thompson K, et al. Clinical and molecular characterization of novel FARS2 variants causing neonatal mitochondrial disease. Mol Genet Metab. 2023;140(3):107657. doi:10.1016/j.ymgme.2023.107657

37. Fine AS, Nemeth CL, Kaufman ML, Fatemi A. Mitochondrial aminoacyl-tRNA synthetase disorders: an emerging group of developmental disorders of myelination. J Neurodev Disord. 2019;11(1):29. doi:10.1186/s11689-019-9292-y

38. Chen X, Liu F, Li B, et al. Neuropathy-associated Fars2 deficiency affects neuronal development and potentiates neuronal apoptosis by impairing mitochondrial function. Cell Biosci. 2022;12(1):103. doi:10.1186/s13578-022-00838-y

39. Liu SS, Jin F, Liu YS, et al. Functional Analysis of the GPI Transamidase Complex by Screening for Amino Acid Mutations in Each Subunit. Molecules. 2021;26(18). doi:10.3390/molecules26185462

40. Murakami Y, Siripanyaphinyo U, Hong Y, Tashima Y, Maeda Y, Kinoshita T. The initial enzyme for glycosylphosphatidylinositol biosynthesis requires PIG-Y, a seventh component. Mol Biol Cell. 2005;16(11):5236–5246. doi:10.1091/mbc.e05-08-0743

41. Daignan-Fornier B, Pinson B. Yeast to Study Human Purine Metabolism Diseases. Cells. 2019;8(1). doi:10.3390/cells8010067

42. Dutto I, Gerhards J, Herrera A, et al. Pathway-specific effects of ADSL deficiency on neurodevelopment. Elife. 2022;11. doi:10.7554/eLife.70518

43. Radenkovic S, Budhraja R, Klein-Gunnewiek T, et al. Neural and metabolic dysregulation in PMM2-deficient human in vitro neural models. Cell Rep. 2024;43(3):113883. doi:10.1016/j.celrep.2024.113883

44. Riley LG, Cowley MJ, Gayevskiy V, et al. A SLC39A8 variant causes manganese deficiency, and glycosylation and mitochondrial disorders. J Inherit Metab Dis. 2017;40(2):261–269. doi:10.1007/s10545-016-0010-6

45. Vetro A, Pisano T, Chiaro S, et al. Early infantile epileptic-dyskinetic encephalopathy due to biallelic PIGP mutations. Neurol Genet. 2020;6(1):e387. doi:10.1212/NXG.0000000000000387

46. Tarailo-Graovac M, Sinclair G, Stockler-Ipsiroglu S, et al. The genotypic and phenotypic spectrum of PIGA deficiency. Orphanet J Rare Dis. 2015;10:23. doi:10.1186/s13023-015-0243-8

47. Cesaroni CA, Contrò G, Spagnoli C, et al. Early-onset dysphagia and severe neurodevelopmental disorder as early signs in a patient with two novel variants in NARS1: a case report and brief review of the literature. Neurogenetics. 2024;25(3):287–291. doi:10.1007/s10048-024-00760-0

48. Menacho C, Okawa S, Alvarez-Merz I, et al. Deep learning-driven neuromorphogenesis screenings identify repurposable drugs for mitochondrial disease. bioRxiv. Published online July 12, 2024. Accessed July 15, 2024. https://www.biorxiv.org/content/10.1101/2024.07.08.602501v1

49. Perlstein EO. Drug repurposing for mitochondrial diseases using a pharmacological model of complex I deficiency in the yeast Yarrowia lipolytica. bioRxiv. Published online January 1, 2020:2020.01.08.899666. doi:10.1101/2020.01.08.899666

50. Singh P. Budding Yeast: An Ideal Backdrop for In vivo Lipid Biochemistry. Front Cell Dev Biol. 2016;4:156. doi:10.3389/fcell.2016.00156

51. Jordá T, Puig S. Regulation of Ergosterol Biosynthesis in Saccharomyces cerevisiae. Genes (Basel). 2020;11(7). doi:10.3390/genes11070795

52. Boonekamp FJ, Knibbe E, Vieira-Lara MA, et al. Full humanization of the glycolytic pathway in Saccharomyces cerevisiae. Cell Rep. 2022;39(13):111010. doi:10.1016/j.celrep.2022.111010

53. Gaynor EC, Mondésert G, Grimme SJ, Reed SI, Orlean P, Emr SD. MCD4 encodes a conserved endoplasmic reticulum membrane protein essential for glycosylphosphatidylinositol anchor synthesis in yeast. Mol Biol Cell. 1999;10(3):627–648. doi:10.1091/mbc.10.3.627

54. Umemura M, Okamoto M, Nakayama K ichi, et al. GWT1 gene is required for inositol acylation of glycosylphosphatidylinositol anchors in yeast. J Biol Chem. 2003;278(26):23639–23647. doi:10.1074/jbc.M301044200

55. Neumüller RA, Gross T, Samsonova AA, et al. Conserved regulators of nucleolar size revealed by global phenotypic analyses. Sci Signal. 2013;6(289):ra70. doi:10.1126/scisignal.2004145

56. Li Z, Vizeacoumar FJ, Bahr S, et al. Systematic exploration of essential yeast gene function with temperature-sensitive mutants. Nat Biotechnol. 2011;29(4):361–367. doi:10.1038/nbt.1832

57. Winzeler EA, Shoemaker DD, Astromoff A, et al. Functional characterization of the S. cerevisiae genome by gene deletion and parallel analysis. Science. 1999;285(5429):901–906. doi:10.1126/science.285.5429.901

58. Galosi S, Edani BH, Martinelli S, et al. De novo DHDDS variants cause a neurodevelopmental and neurodegenerative disorder with myoclonus. Brain. 2022;145(1):208–223. doi:10.1093/brain/awab299

59. Absmanner B, Schmeiser V, Kämpf M, Lehle L. Biochemical characterization, membrane association and identification of amino acids essential for the function of Alg11 from Saccharomyces cerevisiae, an alpha1,2-mannosyltransferase catalysing two sequential glycosylation steps in the formation of the lipid-linked core oligosaccharide. Biochem J. 2010;426(2):205–217. doi:10.1042/BJ20091121

60. Shellhammer JP, Pomeroy AE, Li Y, et al. Quantitative analysis of the yeast pheromone pathway. Yeast. 2019;36(8):495–518. doi:10.1002/yea.3395

